# Knowledge graph analytics platform with LINCS and IDG for Parkinson’s disease target illumination

**DOI:** 10.1101/2020.12.30.424881

**Authors:** Jeremy J Yang, Christopher R Gessner, Joel L Duerksen, Daniel Biber, Jessica L Binder, Murat Ozturk, Brian Foote, Robin McEntire, Kyle Stirling, Ying Ding, David J Wild

## Abstract

**Background:** LINCS, “Library of Integrated Network-based Cellular Signatures”, and IDG, “Illuminating the Druggable Genome”, are both NIH projects and consortia that have generated rich datasets for the study of the molecular basis of human health and disease. LINCS L1000 expression signatures provide unbiased systems/omics experimental evidence. IDG provides compiled and curated knowledge for illumination and prioritization of novel drug target hypotheses. Together, these resources can support a powerful new approach to identifying novel drug targets for complex diseases, such as Parkinson’s disease (PD), which continues to inflict severe harm on human health, and resist traditional research approaches.

**Results:** Integrating LINCS and IDG, we built the Knowledge Graph Analytics Platform (KGAP) to support an important use case: identification and prioritization of drug target hypotheses for associated diseases. The KGAP approach includes strong semantics interpretable by domain scientists and a robust, high performance implementation of a graph database and related analytical methods. Illustrating the value of our approach, we investigated results from queries relevant to PD. Approved PD drug indications from IDG’s resource DrugCentral were used as starting points for evidence paths exploring chemogenomic space via LINCS expression signatures for associated genes, evaluated as target hypotheses by integration with IDG. The KG-analytic scoring function was validated against a gold standard dataset of genes associated with PD as elucidated, published mechanism-of-action drug targets, also from DrugCentral. IDG’s resource TIN-X was used to rank and filter KGAP results for novel PD targets, and one, SYNGR3 (Synaptogyrin-3), was manually investigated further as a case study and plausible new drug target for PD.

**Conclusions:** The synergy of LINCS and IDG, via KG methods, empowers graph analytics methods for the investigation of the molecular basis of complex diseases, and specifically for identification and prioritization of novel drug targets. The KGAP approach enables downstream applications via integration with resources similarly aligned with modern KG methodology. The generality of the approach indicates that KGAP is applicable to many disease areas, in addition to PD, the focus of this paper.

## Background

Integration of heterogeneous datasets is often essential for biomedical knowledge discovery, where relevant evidence may derive from diverse subdomains that include but aren’t limited to: foundational biology, chemistry, clinical data, and social sciences of epidemiology and health economics. Furthermore, there is a need for judicious selection of data types and datasets to integrate, driven by scientific use cases, guided by applicability, accessibility, and veracity of datasets. Accordingly, we have integrated LINCS and IDG to identify and prioritize novel drug target hypotheses, via KG methods and tools. LINCS content includes assay results from cultured and primary human cells treated with bioactive compounds (small molecules or biologics) or genetic perturbations. IDG data utilized in this study include drug-disease associations, gene-disease associations and bibliometric scores from literature text mining.

### Synergy of LINCS and IDG

LINCS and IDG are NIH Common Fund(1) projects, chosen for integration for specific applicability to drug target discovery. LINCS, Library of Networked Cell-based Signatures, is described as a “System-Level Cataloging of Human Cells Response to Perturbations”(2) and features experimental and computational methodology designed to generate useful biomedical knowledge from a systems/omics framework. As a key example, perturbagens (e.g., small molecules, ligands such as growth factors and cytokines, micro-environments, or CRISPR gene over-expression and knockdowns representing disease phenotypes) are characterized by proteomics, transcriptomics (RNA-seq), or biochemical and imaging readouts. Therefore LINCS provides useful mapping from genome to phenome with direct relevance to biomolecular mechanisms and therapeutic hypotheses. IDG(3), with its Target Central Resource Database (TCRD) and data portal Pharos(4), integrates heterogeneous datasets from IDG experimental centers and diverse external sources with a clear purpose, to illuminate understudied (“dark”) protein-coding genes as potential drug targets. IDG is particularly strong in text mining of current biomedical literature and bibliometrics suited for knowledge discovery and evidence evaluation, focused on the sciences of drug discovery research. In addition, another featured resource of IDG is DrugCentral(5). DrugCentral provides information on active ingredients, chemical entities, pharmaceutical products, and their biological targets.

We have combined the complementary strengths of LINCS and IDG to derive new drug discovery insights via these synergies, particularly drug target hypotheses associated with diseases, phenotypes and cell lines. LINCS provides the comprehensive systems/omics view, while IDG focuses on relevant and robust evidence for drug target knowledge and validation.

Table 1 summarizes key concepts and the semantic linkage of LINCS and IDG. Previously, LINCS studies have been designed to identify novel drug targets(6), and IDG has included LINCS as a source for selected data(7). However, our approach provides a new KG method and platform to integrate and add value from these sources for an urgent and unmet scientific use case.

**Table 1:** Key concepts from LINCS and IDG

| LINCS key concepts | IDG key concepts |
| --- | --- |
| **L1000:** Broad Institute microarray platform and LINCS project involving landmark genes.<br>**Landmark Gene:** approximately 1000 genes representing the full human genome, from which a maximum set of gene responses can be inferred.<br>**Cell Lines:** Primary cells and human patient induced pluripotent stem cells (iPSCs) representing diverse tissues and disease states (including ‘healthy’ controls).<br>**Perturbagen:** A perturbagen may be chemical (drug-like), biologics, or genetic.<br>**Signatures:** Various assays (transcriptomics, proteomics, or image analysis) are integrated together to identify patterns of common networks and cellular responses leading to predictive mechanistic interpretations. | **Target:** Drug targets normally defined as human proteins or equivalently protein coding genes.<br>**Mechanism of action:** Known, published, biochemical mechanism of pharmacological effect.<br>**Disease:** Working definition is anything which could be the clinical indication for a drug.<br>**Ligand:** Small molecule with known or hypothesized relationship with one or more targets.<br>**Target Development Level (TDL):** Simple measure of illumination: Tdark, Tbio, Tchem and Tclin.<br>**Target Importance and Novelty:** Bibliometric measures from TIN-X for target prioritization. |
| LINCS-IDG semantic linkage |  |
| LINCS centers utilize in-depth gene and protein expression assays to generate signatures directly mappable to IDG protein targets. Disease and phenotype ontology mapping is a community challenge with useful, working solutions such as UMLS(8). LINCS perturbagens include rigorously defined chemical entities and small molecule drugs included in IDG resource DrugCentral. Therefore, LINCS’s vast number of human cell lines and experimental chemical perturbation datasets, combined with IDG’s protein targets (gene and protein IDs) and DrugCentral active pharmaceutical ingredients (drug compounds) databases, provide a tightly integrated combination resource for drug target discovery. |  |

## Results

### Graph analytics for evidence aggregation

We used the list of PD drugs as starting points for evidence paths in KGAP to identify likely and novel PD targets. KGs and graph analytics provide an intuitive and powerful way to aggregate instances of evidence paths, yielding a score measuring the aggregated evidence. As illustrated schematically in Figure 1, each evidence path connects a drug with expression signature and gene. The magnitude of the expression perturbation is represented by a z-score which can be used for thresholding and/or weighting. The search resulted in 641 genes, ranked by KGAP score produced by the graph analytics algorithm described in Methods and specified by Cypher query Listing 1.

**Figure 1:**
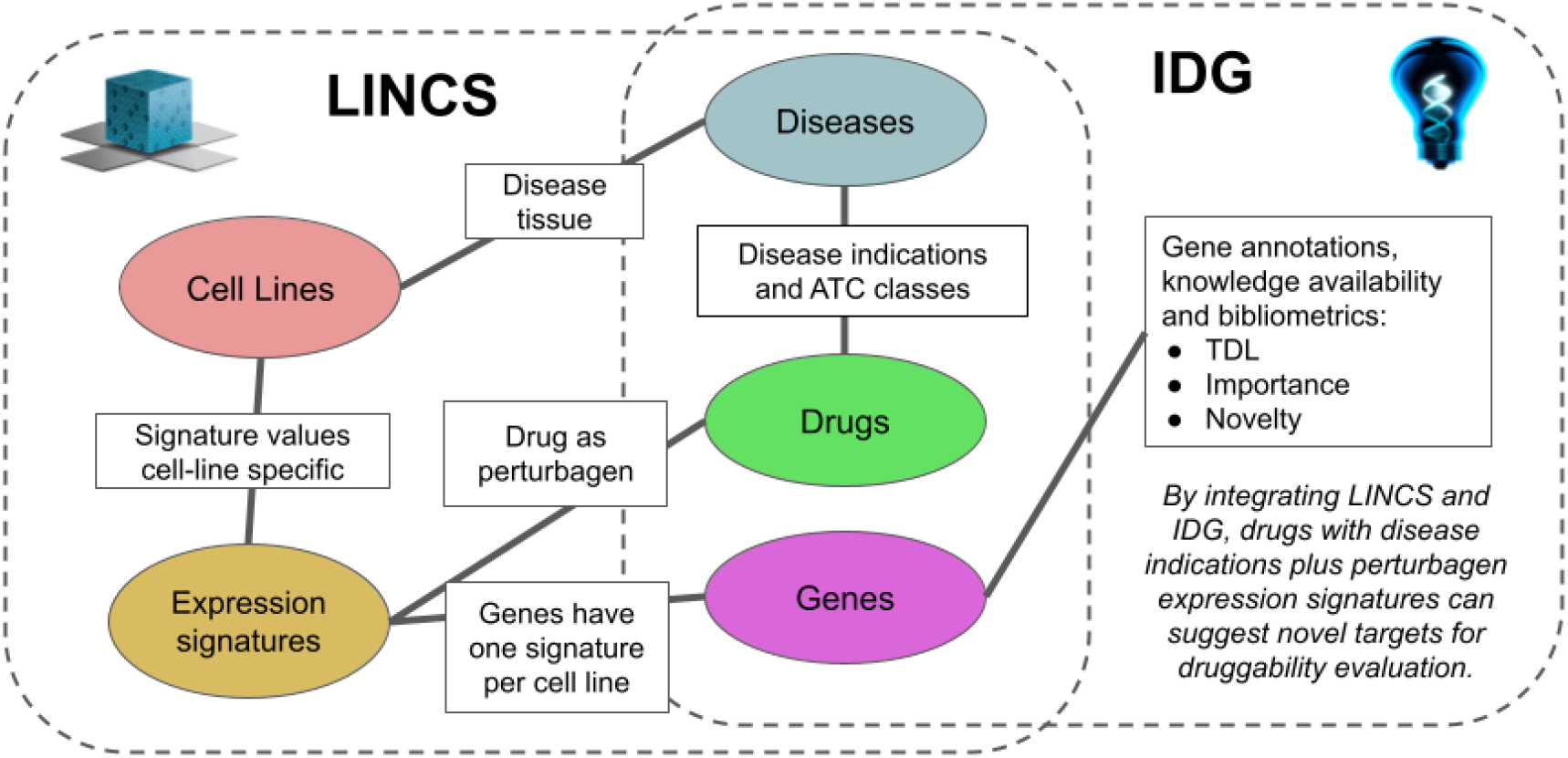
Schematic of overall logic, that strong knowledge of approved drugs and cell lines associate diseases via LINCS expression signatures to differentially expressed genes, for IDG filtering and druggability evaluation.

### Validation versus known mechanism-of-action targets

Solving the fundamental challenges of molecular biomedicine is an ongoing effort. Specifically, there are no gold standard validation datasets of causal or druggable genes for complex diseases, including cancers, neurodegenerative, metabolic, and cardiovascular diseases. Therefore it is pertinent to elucidate the genomic basis of diseases, and the fundamental uncertainties and difficulties concerning the definitions and diagnostic criteria for complex diseases such as PD. Mindful of these difficulties, we validated against a dataset of known drug targets, for the same PD drugs described above, all based on peer-reviewed references, and with elucidated mechanism-of-action (MoA), as manually curated in the DrugCentral database. Given the scientific and regulatory standards for efficacy met by approved drugs, and the standards of evidence for peer-reviewed, manually curated MoA targets, we consider this approach the most useful and informative dataset validation available. LINCS is experimental based, thus derived independently from the curation of MoA targets from DrugCentral, or in machine learning terms, there is strict separation of training and test data.

Figures 2a-d show validation receiver operating characteristic curve (ROC) plots for two variations of our method, and two validation sets. The “D-weighted” variation connotes weighting of evidence paths by the sum of degree at the LINCS expression signatures associating drugs and genes. The “Z-weighted” variation combines degree with the z-score expression level attribute associating signatures and genes. The two validation sets are (1) DrugCentral PD targets, and (2) DrugCentral PD targets with known MoA, as described above. The ROC AUC (area under the curve) values range between 0.64 and 0.74, providing consistent, independent validation of the proposed method for disease to gene association discovery. The validation method, for PD or other disease queries, is reproducible using the source code repository referenced in the Availability of Data and Materials section. Note, the validation presented does not validate KGAP as a classification method. Evaluated as such, given the sparsity of known genes, the specificity is poor. However, classification is not the goal. Rather, our task is to aggregate and assess experimental evidence from LINCS which is applicable to PD as proof of concept. The validation presented simply measures overrepresentation, or enrichment, of known PD genes in relation to KGAP scores, indicating agreement and independent confirmation between two very different sources of knowledge, one purely experimental, the other expertly curated and peer-reviewed publication based.

**Figures 2a-d:**
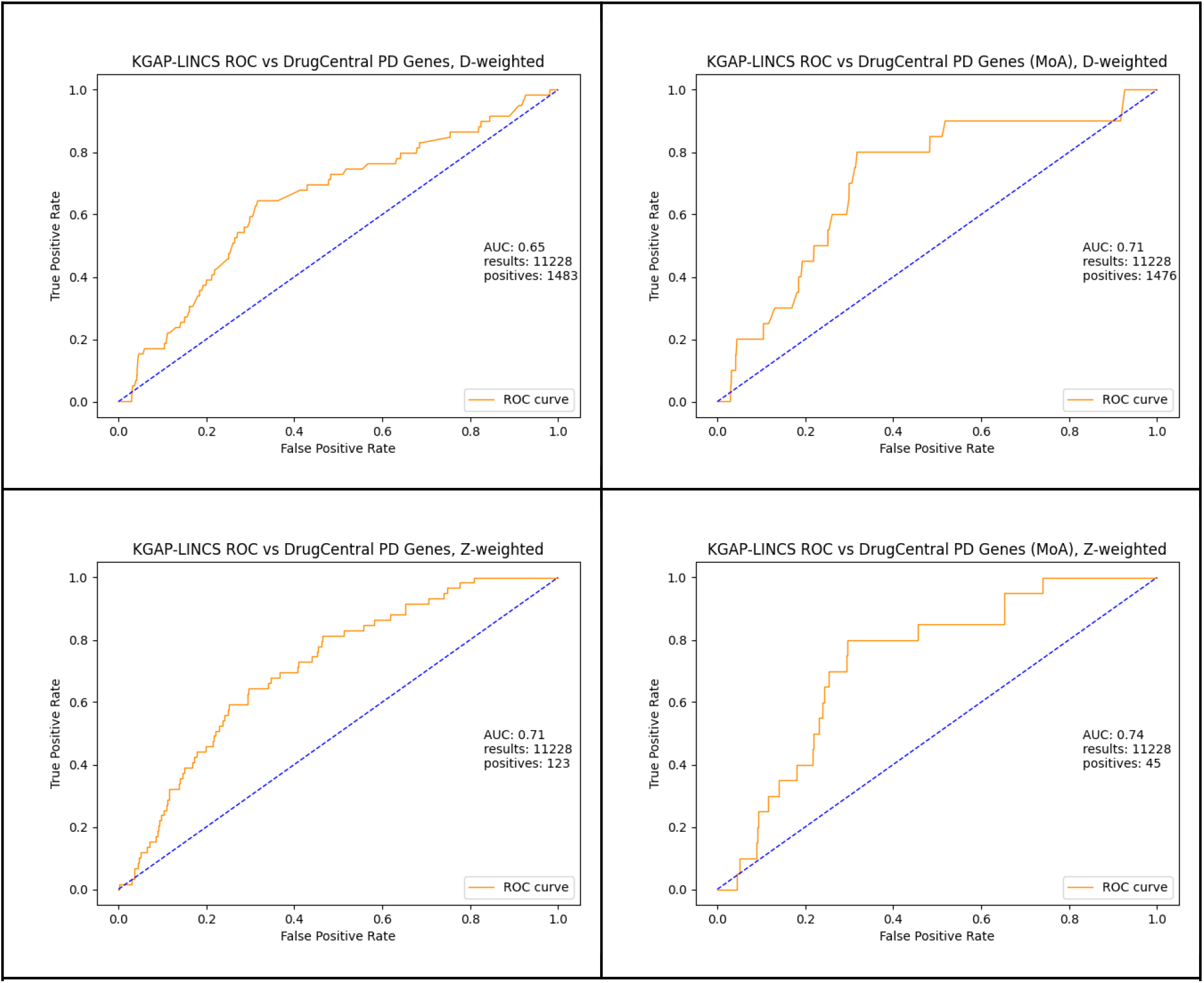
ROC curves with AUC for degree-only and z-score weighted evidence path graph analytics, validated against DrugCentral PD targets and known-with-MoA targets.

### Genes associated with PD, prioritized via IDG

Gene hitlists generated by KGAP were mapped to druggability data from IDG. Target Development Level (TDL) provides a high-level classification into four groups — Tclin, Tchem, Tbio and Tdark — with respect to the depth of investigation from a clinical, chemical and biological standpoint. Further evaluation was explored via Target Importance and Novelty Explorer (TIN-X)(9), an IDG project and bibliometric algorithm for evaluating disease-target associations from scientific literature. Moreover, TIN-X defines ***novelty*** as a bibliometric measure of occurrence rarity in the full PubMed corpus of titles and abstracts, and ***importance*** as a bibliometric measure of co-occurrence associating a specific disease and gene. The premise and motivation for IDG is that many understudied targets could offer new opportunities for medicines of novel therapeutic benefit. Hence TIN-X ranks and presents targets based on the principle of non-dominated solutions optimizing novelty and importance. nds_rank=1 is assigned to all genes relative to which none are superior in both dimensions. nds_rank=2 is this corresponding set with the first set removed, etc. In practice, typical users browse targets beginning at the outer boundary, using TDL color code as an additional guide.

### Case study: SYNGR3, Synaptogyrin-3

To illustrate a typical use case, the KGAP hitlist was browsed for drug target illumination potential based on annotations from IDG. The highest ranked gene classified as Tdark is *SYNGR3*, Synaptogyrin-3. The exact function of SYNGR3 is unclear. However, recently a group provides evidence in the murine brain that *SYNGR3* encodes for a synaptic vesicle protein that interacts with a dopamine transporter(10). One of PD’s hallmark characterizations is the loss of nigrostriatal dopaminergic innervation(11). The high TIN-X rank (nds_rank=6 out of 103) indicates both novelty and importance (PD-relevance), as shown in figure 3a. Additionally, in figure 3b, two reference publications present experimental and theoretical evidence for connection between statin drugs, therapeutic effectiveness for PD, and the gene *SYNGR3*(12,13). Figure 4 displays evidence paths connecting *SYNGR3* with associated expression signatures and drugs, matched by our method.

**Figures 3a-b:**
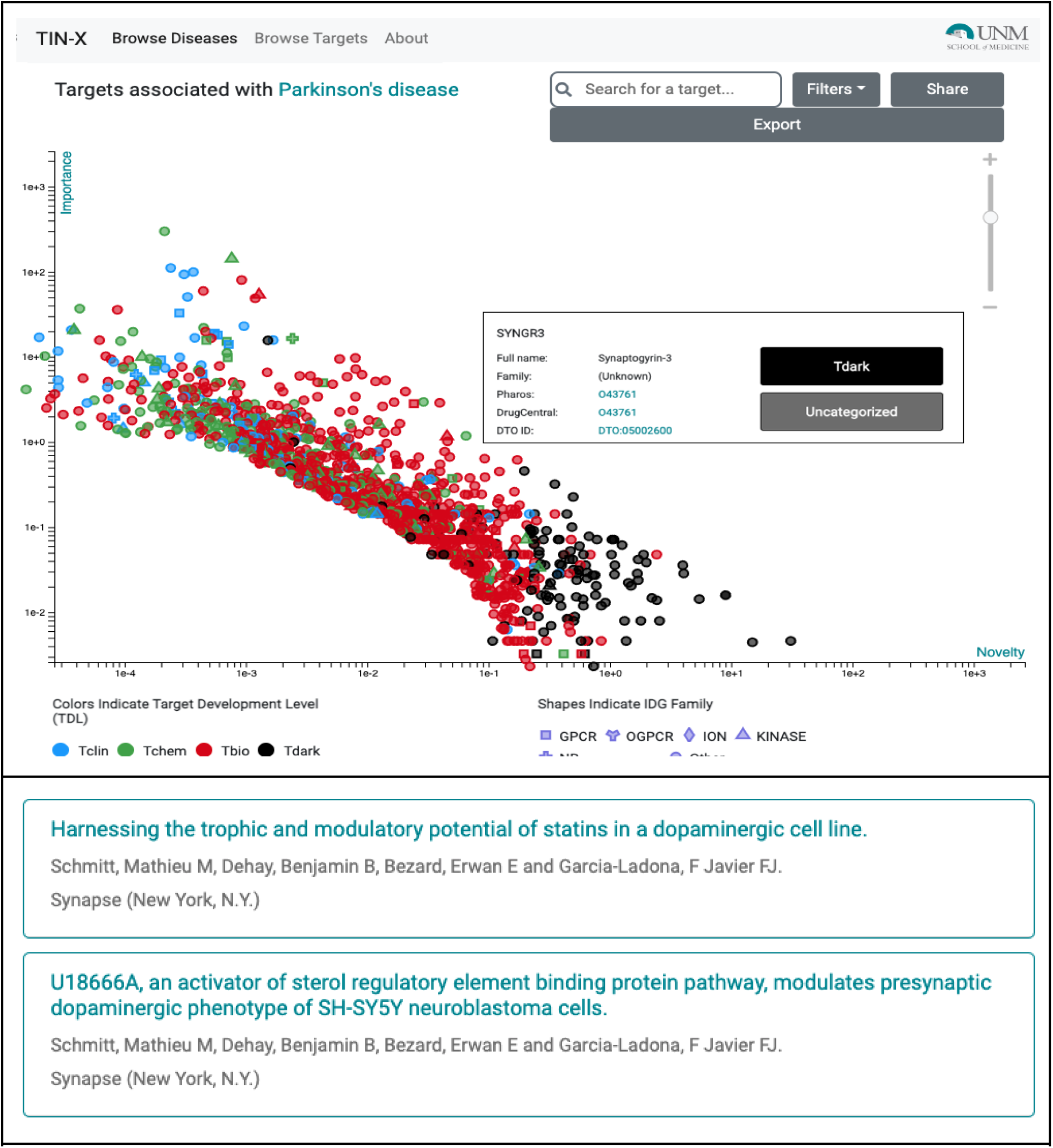
TIN-X scatterplot of genes for Parkinson’s disease, DOID:14330, showing pop-up details for SYNGR3, Synaptogyrin-3, and publication details view for PD associated gene *SYNGR3*.

**Figure 4:**
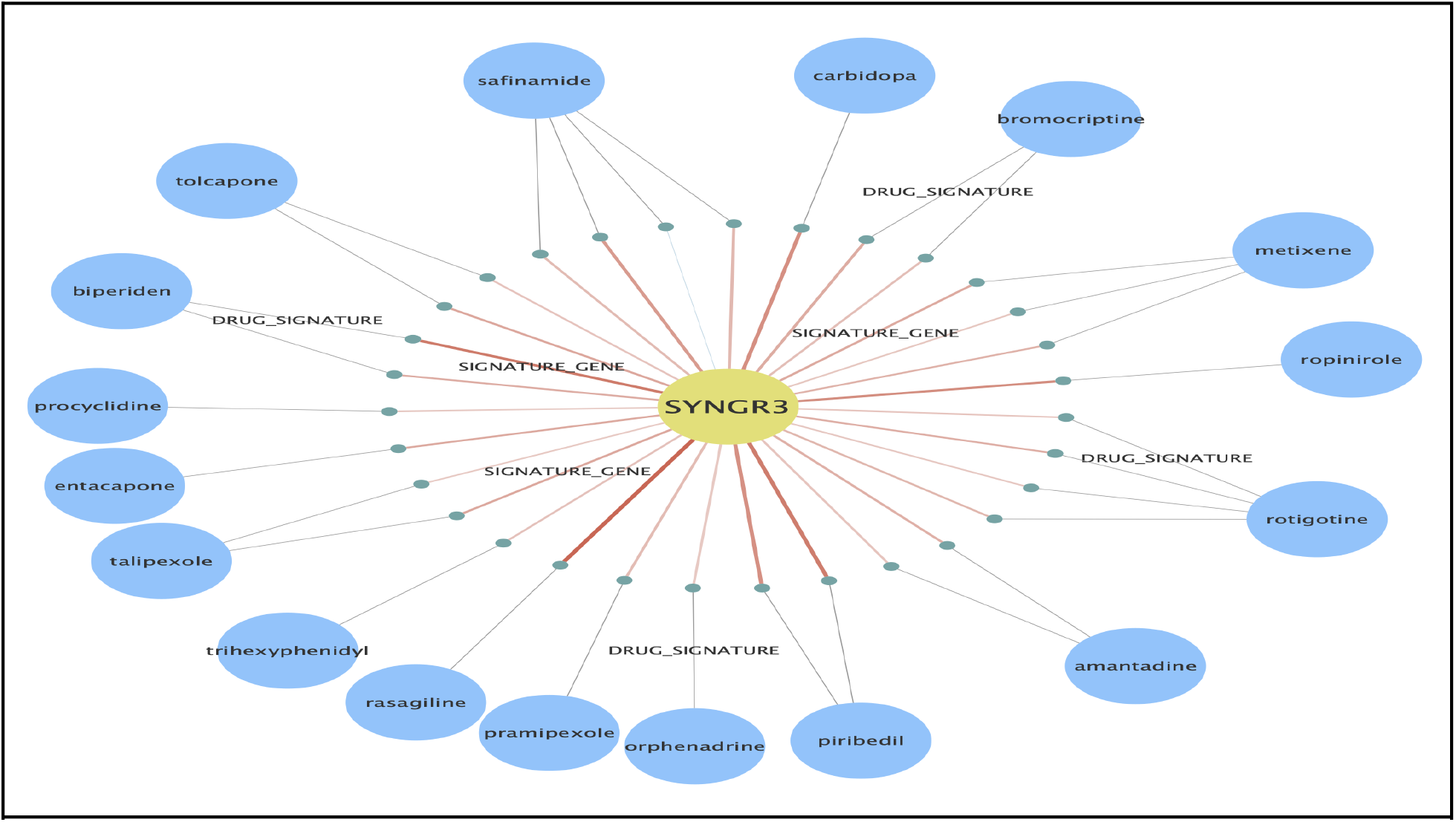
Evidence paths generated from Cypher queries on our Neo4j graph database, for case study *SYNGR3*, showing associated expression signatures and drugs.

## Discussion

This project builds upon numerous prior efforts and resources, conceptual and technological, from domain specific data semantics, to graph database advances, conferring interoperability of depth and breadth(14–24). The contributions cited relate and reflect a prodigious history of multidisciplinary community progress, spanning foundational biomedicine, computer science and informatics, from scientists collaborating effectively and rigorously via computational standards, tools and interfaces. Of particular relevance is the 2015 paper(18) by Himmelstein and colleagues, as it employs a knowledge graph (KG) approach (termed Hetionet) applied to identifying disease-associated genes, closely related to drug target illumination. Other cited work is either not KG-based, or differing in application, and all lack the specificity of our approach both in terms of evidence sources and use-case focus. Another distinctive feature of our approach is its interpretability both algorithmically and in terms of provenance. For example, our validation performance (ROC AUC) is inferior but comparable to the performance reported by Himmelstein (e.g. .74 vs .79), but without training, which introduces knowledge bias, a concern in our domain where ground truth is commonly uncertain or incomplete. Our “machine learning-free” approach relies exclusively on significant LINCS expression signals (z-scores) combined with IDG for target illumination and prioritization.

Regarding the now-common term *knowledge graph (KG)*, there are variations of meaning, but we emphasize: (1) Knowledge with strong semantics via rigorous data modeling; (2) Graph analytics as powerful and intuitive tools for many biomedical applications and users(25). Hence, we use the term KG unambiguously and with specific commitments. Advances in KG technologies have empowered progress far beyond previous efforts. A case in point is Chem2Bio2RDF(14,26), developed in the Wild Lab, enabled by RDF and SPARQL but limited for analytics applications by availability of high-performance triple-store technologies. Another project which one of us (JY) has co-developed is CARLSBAD(27), a bioactivity knowledge graph limited in analytics performance and versatility by its implementation as a relational database. Additional lessons learned relate to (1) Generality versus purposefulness, and (2) Volume versus veracity. Accordingly, this project has prioritized quality and simplicity to mitigate the common problems of (1) unfocused, incoherent interfaces, and (2) unreliable or noisy data. Interpretability also depends on understanding of the tool’s purpose, logic, results, and their provenance and confidence. In many cases, simple and clearly focused tools enhance comprehensibility and usability. In contrast, a toolkit is multi-purpose. For this research the key aim is to find gene associations based on a disease or drug query.

Graph databases are one category of NoSQL (non-SQL or non-relational) databases which are designed to outperform relational databases for some specialized datasets and analyses, and are well suited for representing and exploring complex biological systems(18,28,29). Just as graph diagrams provide an intuitive way to represent and explore relationships between entities, graph databases leverage this intuition to provide both a physical storage method, and approach for combining diverse datasets for exploration and analysis. Neo4j was chosen as an exemplary leader among graph databases with a rich set of graph based capabilities (e.g. Graph Data Science library, and libraries for accessing, flat files, databases, existing open standards such as SPARQL, and open standard in development GQL(30)). In addition, Neo4j offers a free, open source community edition(31), powerful and user-friendly desktop client, APIs, ample documentation, and a vibrant and supportive user community. Though the Neo4j product is featured prominently in this paper, it is important to note that alternative capable graph databases exist, such as DGraph(32) and Amazon Neptune(33). Moreover, Neo4j and other graph databases are built on the shoulders of many diverse contributors and strands of computer science, thus, Neo4j is featured as an exemplar for these advances, and a rational technology choice at this time for implementing our methods.

Development of this knowledge graph analytics platform (KGAP) with LINCS and IDG was motivated by use cases from IDG and its core purpose to illuminate understudied genes and new drug targets. Generally stated: For a given disease, what new drug targets are suggested by the evidence? LINCS data provides aggregated experimental evidence to establish a cell and expression signature based systems biology database, supporting rational aggregation of disease-gene associations, powered by KG analytics well suited both to path-based analytics and interpretation. Representing PD informatically is another key ingredient, addressed by clinical indications for approved drugs, knowledge meeting high regulatory standards. The definitions and nosology of PD and other complex diseases are critical issues and limitations relevant to this study and many others. Any hypotheses regarding PD presumes a useful definition, however biologically PD and its complexity are better described by subtypes and clinical phenotypes. Yet, extant data is generally organized by diseases and clinical diagnoses as historically developed.

## Conclusions

A knowledge graph analytics platform (KGAP) was developed, integrating datasets from LINCS and IDG, for efficient search and aggregation of evidence paths based on a disease query, to identify, score and rank associated genes as drug target hypotheses. Approved indications for prescription drugs were used as high confidence entry points into the KG, and published MoA targets were used as a high confidence validation set. Modern graph database systems such as Neo4j provide a powerful suite of tools for high performance analytics, rigorously and reproducibly encoded, on semantically rich KGs, with interactivity and visualization enhancing human understanding. This study has demonstrated how KGAP can generate novel and plausible drug targets for Parkinson’s disease. It includes a case study of the understudied “Tdark” *SYNGR3* gene, scored highly by KGAP and supported independently by publications identified via the IDG application TIN-X. The KGAP approach and method as implemented is generalizable and applicable to drug target illumination in many other disease areas. Accordingly, in future work we intend to apply KGAP more broadly.

## Methods

### Data Sources

As shown in Figure 1 an understanding of entities and the relationships between them can be quickly represented in a representative graph diagram, this establishes high level concepts and methods for discovering new insights, but also provides a clear design for the required extract transform and load (ETL) steps for building the capability. The datasets listed here were used to construct a graph database instantiation of this design. From LINCS we used the L1000 Phase I and II Level 5 differential gene expression data, also termed the LINCS “Connectivity Map”(34). Level 5 connotes the highest level of processing, normalization, evidence aggregation, and therefore confidence, interpretability, and interoperability. LINCS was processed using the same pipeline used by colleagues for the DrugCentral LINCS drug profiling tool, whereby differential gene expression data across 81 cell-lines were mapped to 1613 unique drug active pharmaceutical ingredients(5). Data and metadata files available from the LINCS Data Portal and NCBI Gene Expression Omnibus (GEO) were ETL-ed via 1.5TB PostgreSQL db. DrugCentral(5) (June 2020 version) was used for indications, ATC codes, chemical structures and other properties. IDG’s TCRD, Target Central Resource Database(35), (v6.7.0) provides gene properties including target development level (TDL: Tclin, Tchem, Tbio, Tdark) and gene family information.

### Data modeling, representing the knowledge

Graphs in Neo4j are composed of nodes and relationships, which correspond with vertices and edges in graph theory terms, respectively. Graph queries can be thought of as patterns for matching paths through the graph, which consist of specified relationship types, each between two nodes, a semantic triple, subject-predicate-object. Neo4j is considered a schema-less database, meaning that data is loaded without constraints. But a strict schema with strong semantics informed by domain knowledge can be implemented, and is essential for accurate knowledge representation and interpretability. The meta-model for a graph can be reported by introspective Cypher query (**`CALL apoc.meta.graph`**). The meta-model for this graph is shown in Figure 5.

**Figure 5:**
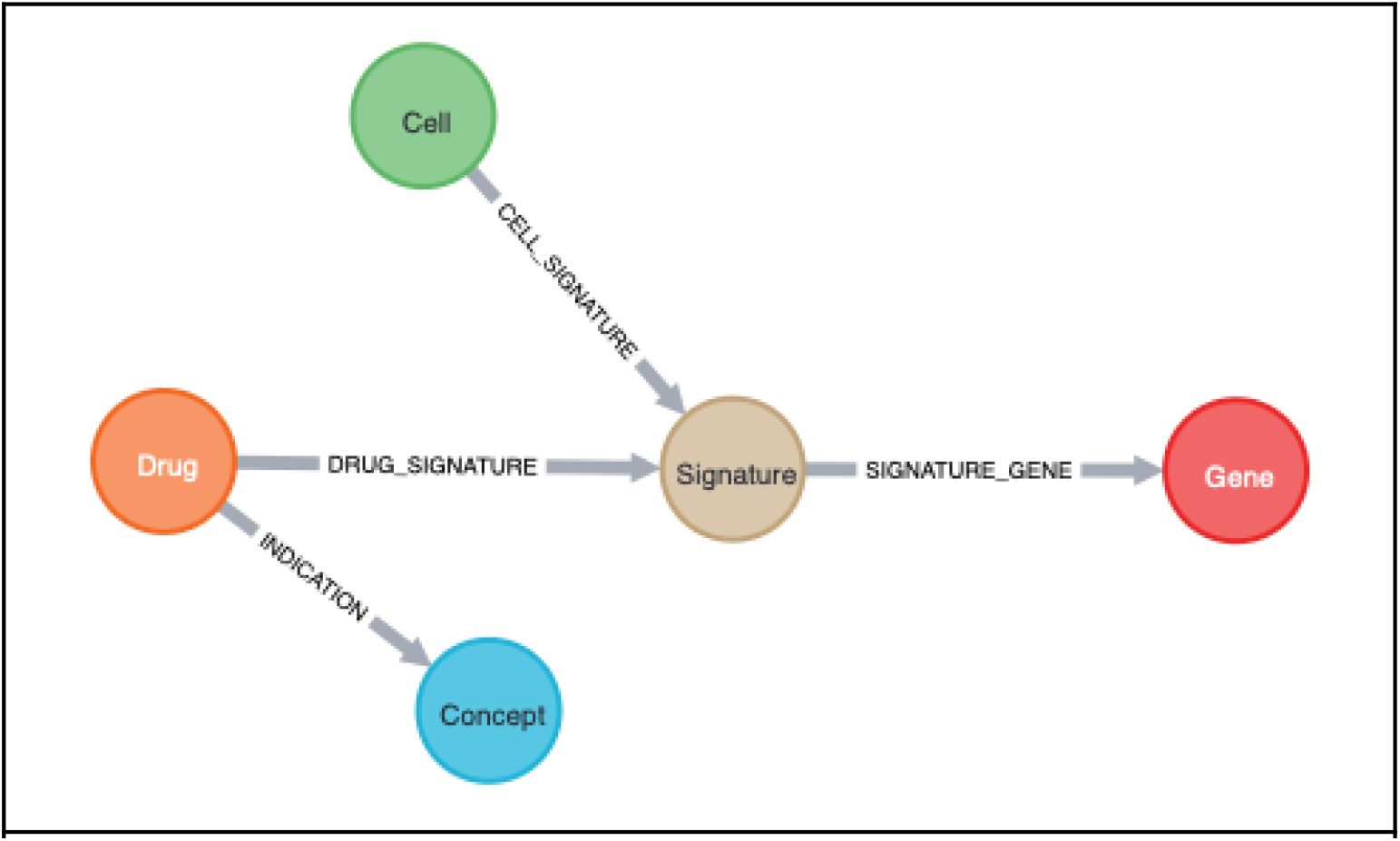
Neo4j meta graph. Both nodes and relationships have properties which can be used in query filtering and analysis.

Our graph is constructed of the following.

#### Relationships

INDICATION is constructed from DrugCentral indications involving a Drug which is also a LINCS perturbagen. CELL_SIGNATURE indicates the Cell type for the LINCS Signature. SIGNATURE_GENE provides the differential expression z-score for the Gene. DRUG_SIGNATURE associates the Drug involved for a given Signature.

#### Nodes

Concept consists of disease terms from DrugCentral with unique OMOP(36) identifiers, and retaining other disease identifiers as properties for interoperability and extensibility. Drug nodes are derived by filtering for LINCS perturbagens which are in DrugCentral and have one or more indications. Genes are from LINCS, with additional properties added from TCRD (e.g., development/druggability level). Cell and Signature nodes are from LINCS without filtering.

Table 2 shows the complete list of relationships present in the graph and count, and Table 3 the list of nodes with counts.

**Table 2:** Type of relationship, source, and total counts

| Vertex type | Edge type | Vertex type | Source | Count |
| --- | --- | --- | --- | --- |
| Signature | SIGNATURE_GENE | Gene | L1000 | 45623637 |
| Cell | CELL_SIGNATURE | Signature | L1000 | 591676 |
| Drug | DRUG_SIGNATURE | Signature | L1000 | 85814 |
| Drug | INDICATION | Concept | DrugCentral, L1000 | 7164 |

**Table 3:**
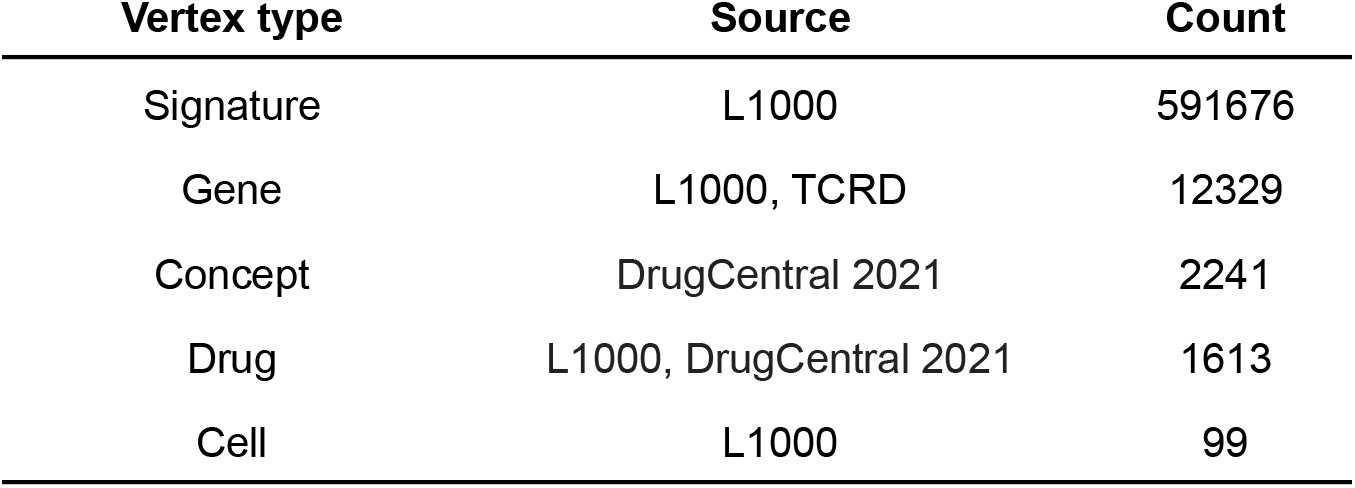
Type of node, primary source/ source of additional properties, and total counts

| Vertex type | Source | Count |
| --- | --- | --- |
| Signature | L1000 | 591676 |
| Gene | L1000, TCRD | 12329 |
| Concept | DrugCentral 2021 | 2241 |
| Drug | L1000, DrugCentral 2021 | 1613 |
| Cell | L1000 | 99 |

We filter the 45.6M signature associations at z-score threshold |z|>3, to identify strong evidence and patterns, which results in 10,663,228 (d:Drug)--(s:Signature)--(g:Gene) paths.

### Parkinson’s disease drug-set

To apply KGAP to PD knowledge discovery, the set of drugs indicated by PD and related conditions were identified using DrugCentral, which derives drug indication from FDA DailyMed (provides trustworthy information about marketed drugs in the United States), and maps SNOMED to OMOP terms(37) for interoperability. A simple substring search for “Parkinson” matches five terms: “Parkinsonism”, “Parkinson’s disease”, “Arteriosclerotic Parkinsonism”, “Dementia associated with Parkinson’s Disease”, and “Neuroleptic-induced Parkinsonism.” Requiring Anatomical and Therapeutic Classification (ATC) “nervous system” is intended to focus on the neurological etiology of PD rather than symptoms. This query returns 25 drugs, shown in Table 4 with their PubChem CID, of which 22 / 25 are present in LINCS. This drug-set represents PD knowledge as in drug-set enrichment analysis (DSEA)(38), analogous to gene-set enrichment analysis (GSEA).

**Table 4:**
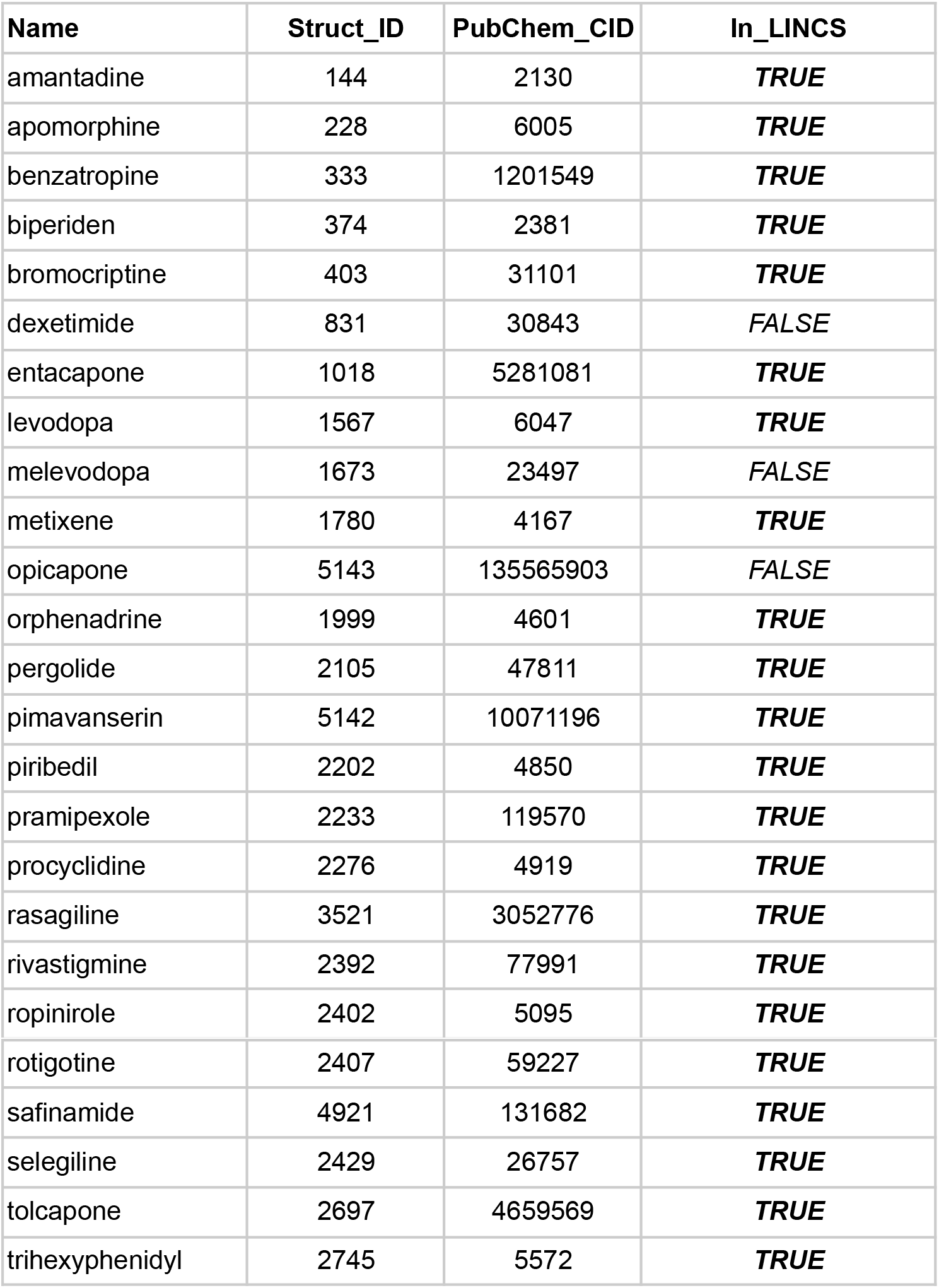
Drug-set representing Parkinson’s disease

| Name | Struct_ID | PubChem_CID | In_LINCS |
| --- | --- | --- | --- |
| amantadine | 144 | 2130 | *TRUE* |
| apomorphine | 228 | 6005 | *TRUE* |
| benzatropine | 333 | 1201549 | *TRUE* |
| biperiden | 374 | 2381 | *TRUE* |
| bromocriptine | 403 | 31101 | *TRUE* |
| dexetimide | 831 | 30843 | *FALSE* |
| entacapone | 1018 | 5281081 | *TRUE* |
| levodopa | 1567 | 6047 | *TRUE* |
| melevodopa | 1673 | 23497 | *FALSE* |
| metixene | 1780 | 4167 | *TRUE* |
| opicapone | 5143 | 135565903 | *FALSE* |
| orphenadrine | 1999 | 4601 | *TRUE* |
| pergolide | 2105 | 47811 | *TRUE* |
| pimavanserin | 5142 | 10071196 | *TRUE* |
| piribedil | 2202 | 4850 | *TRUE* |
| pramipexole | 2233 | 119570 | *TRUE* |
| procyclidine | 2276 | 4919 | *TRUE* |
| rasagiline | 3521 | 3052776 | *TRUE* |
| rivastigmine | 2392 | 77991 | *TRUE* |
| ropinirole | 2402 | 5095 | *TRUE* |
| rotigotine | 2407 | 59227 | **TRUE** |
| safinamide | 4921 | 131682 | **TRUE** |
| selegiline | 2429 | 26757 | **TRUE** |
| tolcapone | 2697 | 4659569 | **TRUE** |
| trihexyphenidyl | 2745 | 5572 | **TRUE** |

### Graph analytics for evidence aggregation

KGAP consists of the graph database described plus code and tools used to query the database for target associations from the input drug-set representing the disease of interest. In the current version of the platform, the queries were implemented via the Neo4j Python Driver(39) and our prototype Python3 command-line applications and notebooks. As in the data modeling in the construction of the database, the Cypher graph queries must be crafted to accurately express the biomedical question, and justify useful interpretation of results. However, since the database was designed precisely for target association and ranking, the Cypher queries involved can be relatively terse. The query used to generate PD target associations is shown in Listing 1. The scoring function is responsible for aggregating evidence from signature counts and z-scores, to reflect that confidence increases with additional confirmatory data, and is an implementation of Stouffer’s function (below) for z-score meta-analysis, based on Fisher’s function using p-values(40). Though not used in KGAP for hypothesis testing, the reference indicates the function is well behaved and combines z-scores as intended.

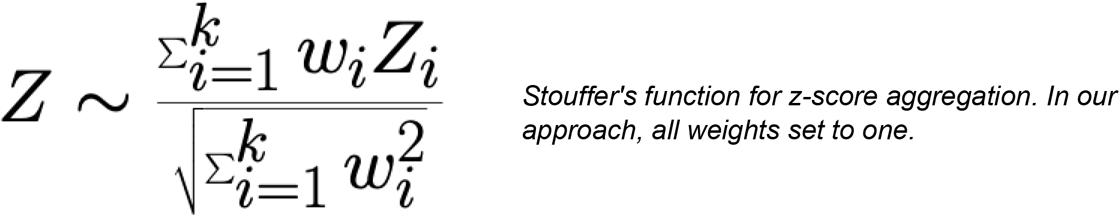

**Listing 1:**
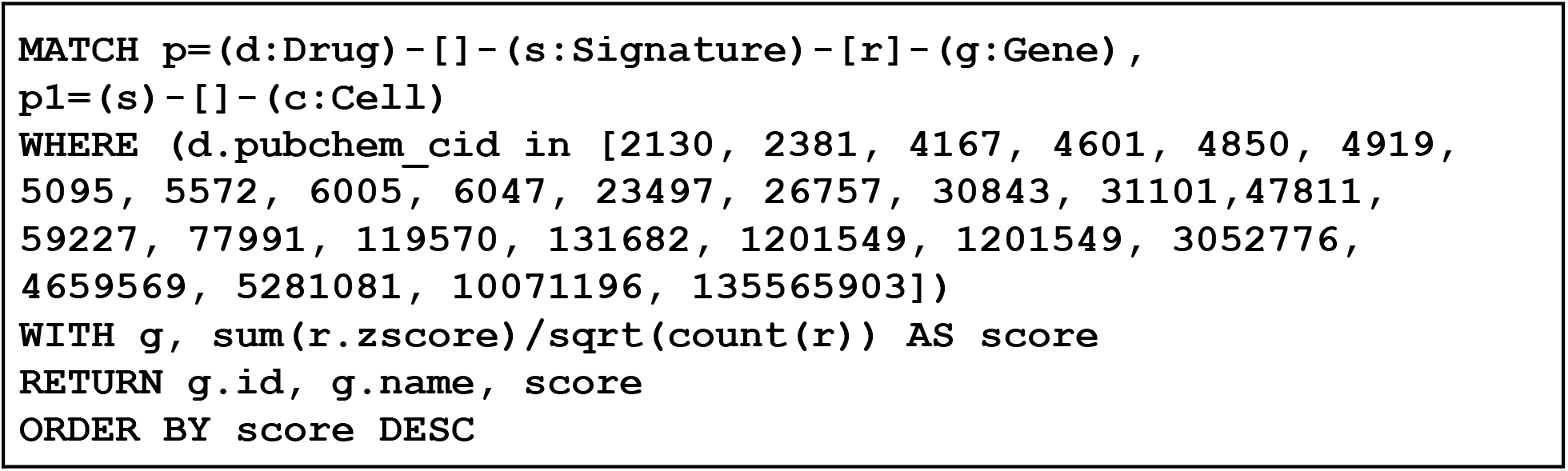
Cypher query used for PD target association.

### Executing the KGAP workflow

In the current version, the workflow is executed via Python3 command-line application **`kgap_anaiysis.py`**. Alternatively, the workflow may be executed via integrated development environment (IDE) such as PyCharm, as shown in Fig. 6. More details, including a Jupyter notebook KGAP workflow implementation, are available via the GitHub repository, https://github.com/IUIDSL/kgap_lincs-idg.

**Figure 6:**
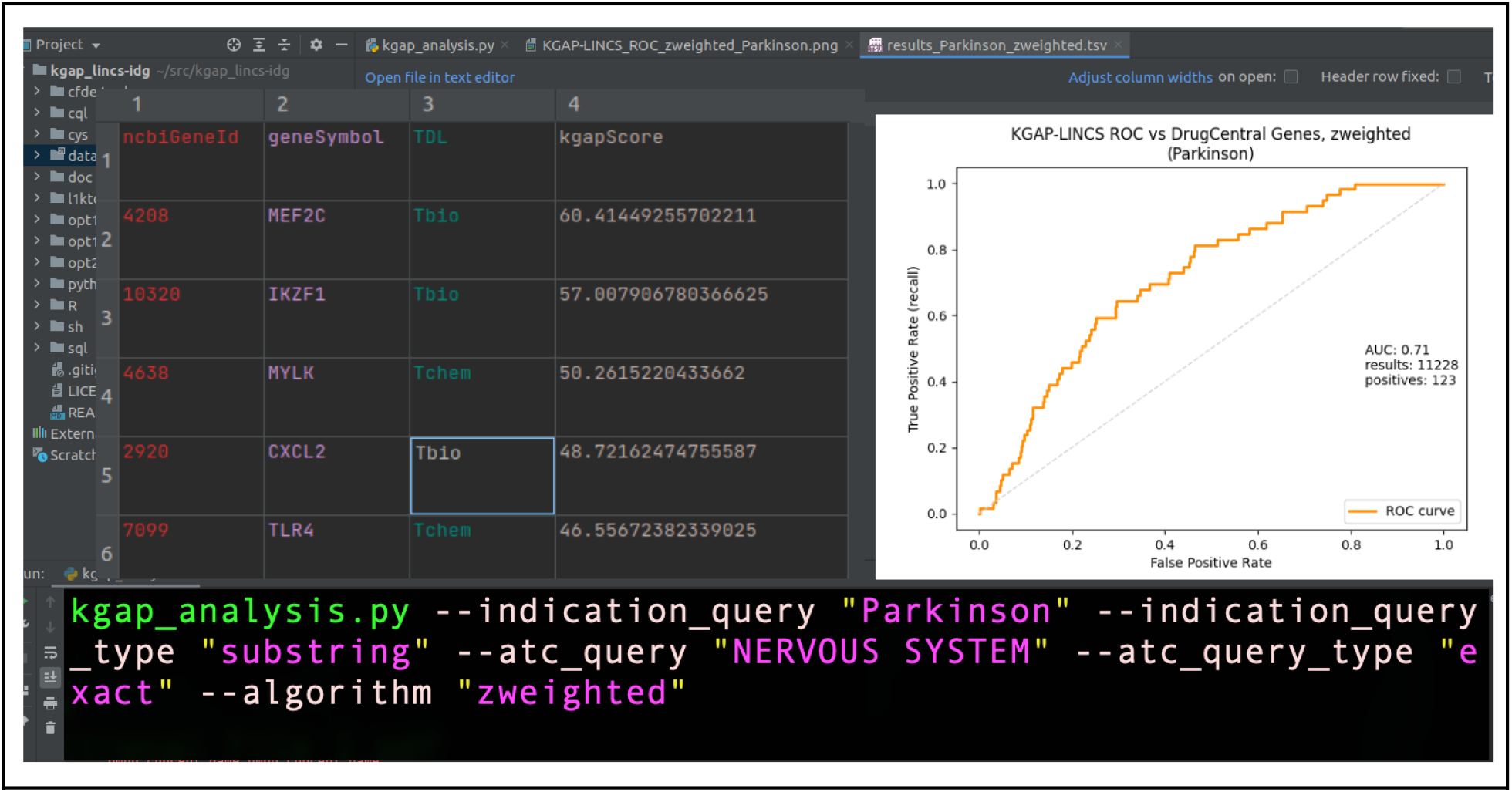
Composite image combining command line executing KGAP for “Parkinson”, output data table, and example ROC plot.

### Exploring and visualizing the KG

Human interaction with the KG through an effective GUI can greatly facilitate understanding and insights. In this project we used (1) Neo4j Desktop, (2) Neo4j Browser web client, used to produce Figure 5, and (3) Cytoscape (v3.8.1) with the Neo4j plugin, which was used to produce Figure 4.

## Supporting information

Tables of results and PD drug-set

## List of abbreviations

ATC: Anatomical Therapeutic Chemical Classification (WHO)
AUROC: Area under receiver operator characteristic (ROC) curve
IDG: Illuminating the druggable genome (NIH Common Fund)
KG: Knowledge graph, also known as a knowledge network
KGAP: Knowledge graph analytics platform
LINCS1000: “Landmark genes” (approximately 1000) from LINCS, chosen for maximal inference of full genomic expression.
LINCS: Library of integrated network-based cellular signatures (NIH Common Fund)
MoA: Mechanism of action, describing biomolecular details of drug effect
PD: Parkinson’s disease
SYNGR3: Synaptogyrin-3, a human gene, subject of case study in this paper
TCRD: Target central resource database (IDG)
TDL: Target development level (IDG)
TIN-X: Target Importance and Novelty Explorer (IDG)

## Declarations

### Ethics approval and consent to participate

Not applicable.

### Consent for publication

Not applicable.

### Availability of data and materials

Source code for processing and analysis is available publicly under Creative Commons Zero v1.0 Universal license at https://github.com/IUIDSL/kgap_lincs-idg. A Knime workflow is employed to extract, transform and load the graph database from sources, and generate TSV files of nodes and relationships. The database is queried, and results reported and visualized using Python and Cypher. A full dump of the KGAP_LINCS-IDG dataset is available at http://cheminfov.informatics.indiana.edu/projects/kgap/data/dclneodb.dump. Source datasets used to build Neo4j database are: (1) DrugCentral-LINCS database dump: http://cheminfov.informatics.indiana.edu/projects/kgap/data/drugcentral_lincs.pgdump; (2) TCRD targets: http://cheminfov.informatics.indiana.edu/projects/kgap/data/tcrd_targets.tsv.gz. Neo4j Community Edition, Python, Scikit-learn, Cytoscape and other packages used in this work are freely and easily findable and accessible.

### Authors’ contributions

JY and DW conceived the project. JD designed and built the graph database and ETL workflows. CG and JD developed graph analytic algorithms. JB led the SYNGR3 case study. JY, JD, DB, CG and MO developed client code. JY, JD, CG, DB, and JB co-authored the manuscript. All reviewed and approved the manuscript.

## Acknowledgements

We are grateful to the LINCS and IDG projects and investigators responsible for the high value datasets shared with the community and used in this research.

