## Supplementary material for "Knowledge graph analytics platform with LINCS and IDG for Parkinson’s disease target illumination": Tables of results and PD drug-set

### Additional File 1

Supplementary materials **2**
Top scoring 100 genes associated with Parkinson's disease 2
Drugs indicated for Parkinson's disease and with ATC "NERVOUS SYSTEM" with structures and properties 6

### Supplementary materials

#### Top scoring 100 genes associated with Parkinson's disease

Full results with all associated genes are provided as Additional File 2 (KGAP_gene_results.tsv) and Additional File 3 (KGAP_gene_results.xlsx).

| | Symbol | Family | Name | KGAP Score | IDG TDL | TINX nds_rank | TINX Importance | TINX Novelty |
|---|---|---|---|---|---|---|---|---|
| 1 | MYLK | Kinase | Myosin light chain kinase, smooth muscle | 50.2615 | Tchem | 77 | 0.0385 | 0.0013 |
| 2 | CXCL2 | | C-X-C motif chemokine 2 | 48.7216 | Tbio | 100 | 0.0100 | 0.0030 |
| 3 | SNAP25 | | Synaptosomal-associated protein 25 | 46.0216 | Tclin | 21 | 0.5410 | 0.0021 |
| 4 | SPP1 | | Osteopontin | 45.5870 | Tbio | 38 | 0.6174 | 0.0002 |
| 5 | PAK1 | Kinase | Serine/threonine-protein kinase PAK 1 | 44.6944 | Tchem | 80 | 0.0286 | 0.0030 |
| 6 | SRC | Kinase | Proto-oncogene tyrosine-protein kinase Src | 43.9768 | Tclin | 21 | 1.4295 | 0.0001 |
| 7 | IDE | Enzyme | Insulin-degrading enzyme | 42.9433 | Tchem | 55 | 0.1012 | 0.0025 |
| 8 | ST3GAL5 | Enzyme | Lactosylceramide alpha-2,3-sialyltransferase | 42.2349 | Tbio | 40 | 0.0455 | 0.0110 |
| 9 | NR3C1 | NR | Glucocorticoid receptor | 41.6000 | Tclin | 30 | 0.8131 | 0.0003 |
| 10 | MAP7 | | Ensconsin | 38.3868 | Tbio | 38 | 0.0204 | 0.0274 |
| 11 | HBB | | Hemoglobin subunit beta | 37.9577 | Tbio | 42 | 0.4245 | 0.0004 |
| 12 | BMP4 | | Bone morphogenetic protein 4 | 37.9529 | Tchem | 47 | 0.2492 | 0.0006 |
| 13 | ADRB2 | GPCR | Beta-2 adrenergic receptor | 37.0831 | Tclin | 28 | 0.9421 | 0.0004 |
| 14 | SLC1A4 | Transporter | Neutral amino acid transporter A | 36.8827 | Tbio | 44 | 0.0286 | 0.0164 |
| 15 | SLC1A4 | Transporter | Neutral amino acid transporter A | 36.8827 | Tbio | 44 | 0.0286 | 0.0164 |
| 16 | NOTCH1 | | Neurogenic locus notch homolog protein 1 | 36.5929 | Tchem | 30 | 0.9557 | 0.0001 |
| 17 | MAMLD1 | | Mastermind-like domain-containing protein 1 | 36.3975 | Tbio | 34 | 0.1000 | 0.0095 |

| | | | | | | | | |
|---|---|---|---|---|---|---|---|---|
| 18 | GPER1 | GPCR | G-protein coupled estrogen receptor 1 | 35.8607 | Tchem | 21 | 0.6166 | 0.0015 |
| 19 | MCOLN1 | IC | Mucolipin-1 | 35.2058 | Tchem | 58 | 0.0788 | 0.0028 |
| 20 | PTGS2 | Enzyme | Prostaglandin G/H synthase 2 | 34.9171 | Tclin | 10 | 8.6930 | 0.0001 |
| 21 | PTGS2 | Enzyme | Prostaglandin G/H synthase 2 | 34.9171 | Tclin | 10 | 8.6930 | 0.0001 |
| 22 | SLC25A14 | Transporter | Brain mitochondrial carrier protein 1 | 34.6732 | Tbio | 17 | 0.1250 | 0.0313 |
| 23 | EPHB2 | Kinase | Ephrin type-B receptor 2 | 34.5689 | Tchem | 24 | 0.5397 | 0.0019 |
| 24 | SIRT3 | Epigenetic | NAD-dependent protein deacetylase sirtuin-3, mitochondrial | 34.2580 | Tchem | 16 | 1.1718 | 0.0013 |
| 25 | IL1B | | Interleukin-1 beta | 32.9557 | Tclin | 5 | 20.7147 | 0.0000 |
| 26 | STAP2 | | Signal-transducing adaptor protein 2 | 32.8113 | Tbio | 53 | 0.0370 | 0.0061 |
| 27 | AGL | | Glycogen debranching enzyme | 31.0804 | Tbio | 66 | 0.0588 | 0.0024 |
| 28 | NFATC3 | TF | Nuclear factor of activated T-cells, cytoplasmic 3 | 30.8690 | Tbio | 36 | 0.1111 | 0.0076 |
| 29 | PROS1 | | Vitamin K-dependent protein S | 30.8634 | Tbio | 61 | 0.1052 | 0.0010 |
| 30 | ADGRE5 | GPCR | CD97 antigen | 30.2677 | Tbio | 52 | 0.0159 | 0.0132 |
| 31 | ADO | Enzyme | 2-aminoethanethiol dioxygenase | 30.0236 | Tbio | 49 | 0.1429 | 0.0019 |
| 32 | CNOT4 | | CCR4-NOT transcription complex subunit 4 | 29.6965 | Tbio | 28 | 0.3537 | 0.0021 |
| 33 | IKBKE | Kinase | Inhibitor of nuclear factor kappa-B kinase subunit epsilon | 29.4758 | Tchem | 48 | 0.0139 | 0.0184 |
| 34 | DFFB | | DNA fragmentation factor subunit beta | 28.8786 | Tbio | 71 | 0.0083 | 0.0062 |
| 35 | RELB | TF | Transcription factor RelB | 28.5684 | Tbio | 58 | 0.0714 | 0.0032 |
| 36 | CDK6 | Kinase | Cyclin-dependent kinase 6 | 28.5364 | Tclin | 58 | 0.1052 | 0.0017 |
| 37 | TIAM1 | | T-lymphoma invasion and metastasis-inducing protein 1 | 27.6093 | Tbio | 59 | 0.0357 | 0.0053 |
| 38 | SYNE2 | | Nesprin-2 | 27.5690 | Tbio | 108 | 0.0044 | 0.0019 |

| | | | | | | | | |
|---|---|---|---|---|---|---|---|---|
| 39 | MAP2K5 | Kinase | Dual specificity mitogen-activated protein kinase kinase 5 | 27.5558 | Tchem | 46 | 0.0385 | 0.0091 |
| 40 | ADAM10 | Enzyme | Disintegrin and metalloproteinase domain-containing protein 10 | 27.4248 | Tchem | 61 | 0.0893 | 0.0021 |
| 41 | LYN | Kinase | Tyrosine-protein kinase Lyn | 27.3761 | Tclin | 49 | 0.1447 | 0.0014 |
| 42 | RAC2 | Enzyme | Ras-related C3 botulinum toxin substrate 2 | 26.4949 | Tbio | 73 | 0.0635 | 0.0012 |
| 43 | HMOX1 | Enzyme | Heme oxygenase 1 | 26.3646 | Tchem | 8 | 9.5207 | 0.0002 |
| 44 | HLA-DRA | | HLA class II histocompatibility antigen, DR alpha chain | 26.2979 | Tbio | 7 | 1.1049 | 0.0071 |
| 45 | CASP2 | Enzyme | Caspase-2 | 26.2581 | Tchem | 25 | 0.4140 | 0.0022 |
| 46 | STAT3 | TF | Signal transducer and activator of transcription 3 | 25.7421 | Tchem | 22 | 1.3099 | 0.0001 |
| 47 | CXCR4 | GPCR | C-X-C chemokine receptor type 4 | 25.3096 | Tclin | 49 | 0.2897 | 0.0003 |
| 48 | CXCR4 | GPCR | C-X-C chemokine receptor type 4 | 25.3096 | Tclin | 49 | 0.2897 | 0.0003 |
| 49 | GNA15 | | Guanine nucleotide-binding protein subunit alpha-15 | 24.9719 | Tbio | 69 | 0.0680 | 0.0018 |
| 50 | HP | | Haptoglobin | 24.7161 | Tbio | 39 | 0.5516 | 0.0002 |
| 51 | SYNGR3 | | Synaptogyrin-3 | 24.2133 | Tdark | 6 | 0.0524 | 0.2604 |
| 52 | SOX2 | TF | Transcription factor SOX-2 | 24.0188 | Tbio | 24 | 1.0835 | 0.0005 |
| 53 | ETV1 | TF | ETS translocation variant 1 | 23.8861 | Tbio | 34 | 0.1429 | 0.0070 |
| 54 | MNAT1 | Enzyme | CDK-activating kinase assembly factor MAT1 | 23.8825 | Tbio | 51 | 0.0417 | 0.0072 |
| 55 | MNAT1 | Enzyme | CDK-activating kinase assembly factor MAT1 | 23.8825 | Tbio | 51 | 0.0417 | 0.0072 |
| 56 | PRKAG2 | Enzyme | 5'-AMP-activated protein kinase subunit gamma-2 | 23.8621 | Tbio | 28 | 0.1429 | 0.0094 |
| 57 | RRP8 | Enzyme | Ribosomal RNA-processing protein 8 | 23.6185 | Tbio | 18 | 0.3571 | 0.0046 |
| 58 | DAXX | | Death domain-associated protein 6 | 23.3585 | Tbio | 24 | 0.2625 | 0.0043 |
| 59 | CES1 | Enzyme | Liver carboxylesterase 1 | 22.8917 | Tchem | 58 | 0.1119 | 0.0013 |

| | | | | | | | | |
|---|---|---|---|---|---|---|---|---|
| 60 | PIK3R3 | Enzyme | Phosphatidylinositol 3-kinase regulatory subunit gamma | 22.8118 | Tbio | 26 | 0.0714 | 0.0222 |
| 61 | KIT | Kinase | Mast/stem cell growth factor receptor Kit | 22.6919 | Tclin | 54 | 0.2085 | 0.0001 |
| 62 | EGF | | Pro-epidermal growth factor | 22.4146 | Tbio | 19 | 1.5146 | 0.0002 |
| 63 | LPL | Enzyme | Lipoprotein lipase | 22.2644 | Tchem | 77 | 0.0623 | 0.0007 |
| 64 | LPL | Enzyme | Lipoprotein lipase | 22.2644 | Tchem | 77 | 0.0623 | 0.0007 |
| 65 | STAT5B | TF | Signal transducer and activator of transcription 5B | 21.8759 | Tchem | 76 | 0.0658 | 0.0009 |
| 66 | GATA3 | TF | Trans-acting T-cell-specific transcription factor GATA-3 | 21.2262 | Tbio | 88 | 0.0337 | 0.0009 |
| 67 | CAPN1 | | Calpain-1 catalytic subunit | 21.0522 | Tchem | 24 | 0.4176 | 0.0024 |
| 68 | ERBB3 | Kinase | Receptor tyrosine-protein kinase erbB-3 | 21.0286 | Tclin | 47 | 0.2109 | 0.0010 |
| 69 | IKBKB | Kinase | Inhibitor of nuclear factor kappa-B kinase subunit beta | 20.6495 | Tchem | 26 | 0.6194 | 0.0011 |
| 70 | SYK | Kinase | Tyrosine-protein kinase SYK | 20.5050 | Tclin | 74 | 0.0748 | 0.0007 |
| 71 | NTS | | Neurotensin/neuromedin N | 20.1709 | Tbio | 10 | 4.1146 | 0.0004 |
| 72 | CLPX | Enzyme | ATP-dependent Clp protease ATP-binding subunit clpX-like, mitochondrial | 20.1206 | Tbio | 67 | 0.0227 | 0.0046 |
| 73 | ACLY | Enzyme | ATP-citrate synthase | 20.1079 | Tclin | 110 | 0.0058 | 0.0014 |
| 74 | TF | | Serotransferrin | 20.0698 | Tbio | 61 | 0.0786 | 0.0023 |
| 75 | LTF | | Lactotransferrin | 19.9048 | Tbio | 33 | 0.3492 | 0.0014 |
| 76 | CALM3 | | Calmodulin-3 | 19.8514 | Tclin | 57 | 0.0185 | 0.0083 |
| 77 | ALDOC | Enzyme | Fructose-bisphosphate aldolase C | 19.8028 | Tbio | 21 | 0.1170 | 0.0214 |
| 78 | CEBPA | TF | CCAAT/enhancer-binding protein alpha | 19.7968 | Tbio | 90 | 0.0238 | 0.0010 |
| 79 | SOCS2 | | Suppressor of cytokine signaling 2 | 19.3824 | Tbio | 61 | 0.0292 | 0.0046 |
| 80 | LCN2 | Enzyme | Neutrophil gelatinase-associated lipocalin | 19.3678 | Tbio | 53 | 0.2036 | 0.0004 |
| 81 | GATA2 | TF | Endothelial transcription factor GATA-2 | 19.2315 | Tbio | 32 | 0.3113 | 0.0018 |
| 82 | PTPRC | Enzyme | Receptor-type tyrosine-protein phosphatase C | 19.0457 | Tchem | 28 | 0.9965 | 0.0002 |

| | | | | | | | | |
|---|---|---|---|---|---|---|---|---|
| 83 | DDR1 | Kinase | Epithelial discoidin domain-containing receptor 1 | 18.8839 | Tchem | 51 | 0.0660 | 0.0048 |
| 84 | EIF4G1 | | Eukaryotic translation initiation factor 4 gamma 1 | 18.6275 | Tbio | 7 | 2.4872 | 0.0022 |
| 85 | RBP4 | | Retinol-binding protein 4 | 18.5690 | Tchem | 59 | 0.0976 | 0.0019 |
| 86 | KYNU | Enzyme | Kynureninase | 18.5648 | Tchem | 32 | 0.0373 | 0.0241 |
| 87 | CDK5R 1 | Enzyme | Cyclin-dependent kinase 5 activator 1 | 18.5556 | Tchem | 21 | 0.1447 | 0.0158 |
| 88 | SELL | | L-selectin | 18.2829 | Tchem | 88 | 0.0357 | 0.0004 |
| 89 | PPBP | | Platelet basic protein | 18.0505 | Tbio | 91 | 0.0286 | 0.0006 |
| 90 | CTTN | | Src substrate cortactin | 17.7222 | Tbio | 61 | 0.0847 | 0.0021 |
| 91 | FOXO4 | TF | Forkhead box protein O4 | 17.3937 | Tbio | 60 | 0.0476 | 0.0032 |
| 92 | FOXO4 | TF | Forkhead box protein O4 | 17.3937 | Tbio | 60 | 0.0476 | 0.0032 |
| 93 | BCL2 | | Apoptosis regulator Bcl-2 | 17.2447 | Tclin | 24 | 0.2166 | 0.0057 |
| 94 | RAB27 A | Enzyme | Ras-related protein Rab-27A | 17.2295 | Tchem | 39 | 0.1296 | 0.0046 |
| 95 | CADM1 | | Cell adhesion molecule 1 | 17.2064 | Tbio | 109 | 0.0071 | 0.0013 |
| 96 | TXNIP | | Thioredoxin-interacting protein | 17.1377 | Tbio | 33 | 0.2018 | 0.0028 |
| 97 | SORBS 3 | | Vinexin | 17.0079 | Tbio | 28 | 0.0590 | 0.0250 |
| 98 | NTRK2 | Kinase | BDNF/NT-3 growth factors receptor | 16.9698 | Tclin | 9 | 2.8107 | 0.0006 |
| 99 | SERPIN A1 | | Alpha-1-antitrypsin | 16.8725 | Tbio | 39 | 0.5851 | 0.0002 |
| 100 | PLCB3 | Enzyme | 1-phosphatidylinositol 4,5-bisphosphate phosphodiesterase beta-3 | 16.7498 | Tbio | 59 | 0.0084 | 0.0117 |

#### Drugs indicated for Parkinson's disease and with ATC "NERVOUS SYSTEM" with structures and properties

Also available for convenience and interoperability as Additional File 4 (PD_drugs.tsv) and Additional File 5 (PD_drugs.xlsx).

| Name | DrugCent | PubChem | Molecular | SMILES | InChIKey | ATC level1 | IN_LINC |
|---|---|---|---|---|---|---|---|

| | ral struct_id | CID | Depiction | | | class | S |
|---|---|---|---|---|---|---|---|
| amantadine | 144 | 2130 | 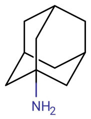 | NC12CC3CC(CC(C3)C1)C2 | DKNWSYNQZKUICI-UHFFFAOYSA-N | NERVOUS SYSTEM | ***TRUE*** |
| apomorphine | 228 | 6005 | 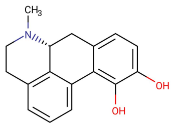 | CN1CCC2=C3[C@H]1CC1=CC=C(O)C(O)=C1C3=CC=C2 | VMWNQDUVQKEIOC-CYBMUJFWSA-N | NERVOUS SYSTEM | ***TRUE*** |
| benzatropine | 333 | 1201549 | 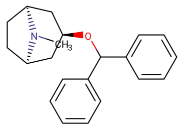 | CN1[C@H]2CC[C@@H]1C[C@@H](C2)OC(C1=CC=CC=C1)C1=CC=CC=C1 | GIJXKZJWITVLHI-PMOLBWCYSA-N | NERVOUS SYSTEM | ***TRUE*** |
| biperiden | 374 | 2381 | 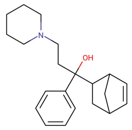 | OC(CCN1CCCCC1)(C1CC2CC1C=C2)C1=CC=CC=C1 | YSXKPIUOCJLQIE-UHFFFAOYSA-N | NERVOUS SYSTEM | ***TRUE*** |
| bromocriptine | 403 | 31101 | 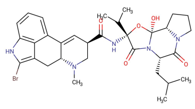 | CC(C)C[C@@H]1N2C(=O)[C@](NC(=O)[C@H]3CN(C)[C@@H]4CC5=C(Br)NC6=C5C(=CC=C6)C4=C3)(O[C@@]2(O)[C@@H]2CCCN2C1=O)C(C)C | OZVBMTJYIDMWIL-AYFBDAFISA-N | NERVOUS SYSTEM | ***TRUE*** |

| | | | | | | | |
|---|---|---|---|---|---|---|---|
| dexetimide | 831 | 30843 | 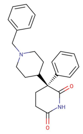 | O=C1CC[C@@](C2CCN(CC3=CC=CC=C3)CC2)(C(=O)N1)C1=CC=CC=C1 | LQQIVYSCPWCSSD-HSZRJFAPSA-N | NERVOUS SYSTEM | *FALSE* |
| entacapone | 1018 | 5281081 | 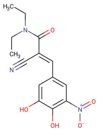 | CCN(CC)C(=O)C(=C\C1=CC(=C(O)C(O)=C1)[N+]([O-])=O)\C#N | JRURYQJSLYLRLN-BJMVGYQFSA-N | NERVOUS SYSTEM | ***TRUE*** |
| levodopa | 1567 | 6047 | 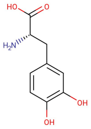 | N[C@@H](CC1=CC(O)=C(O)C=C1)C(O)=O | WTDRDQBEARUVNC-LURJTMIESA-N | NERVOUS SYSTEM | ***TRUE*** |
| melevodopa | 1673 | 23497 | 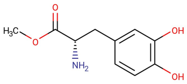 | COC(=O)[C@@H](N)CC1=CC(O)=C(O)C=C1 | XBBDACCLCFWBSI-ZETCQYMHSA-N | NERVOUS SYSTEM | *FALSE* |
| metixene | 1780 | 4167 | 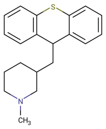 | CN1CCCC(CC2C3=CC=CC=C3SC3=C2C=CC=C3)C1 | MJFJKKXQDNNUJF-UHFFFAOYSA-N | NERVOUS SYSTEM | ***TRUE*** |
| opicapone | 5143 | 135565903 | 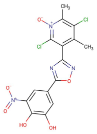 | CC1=C(C2=NOC(=N2)C2=CC(O)=C(O)C(=C2)[N+]([O-])=O)C(Cl)=[N+]([O-])C(C)=C1Cl | ASOADIZOVZTJSR-UHFFFAOYSA-N | NERVOUS SYSTEM | *FALSE* |
| orphenadrine | 1999 | 4601 | 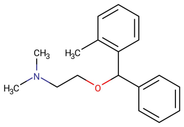 | CN(C)CCOC(C1=CC=CC=C1)C1=C(C)C=CC=C1 | QVYRGXJJSLMXQH-UHFFFAOYSA-N | NERVOUS SYSTEM | ***TRUE*** |

| | | | | | | | |
|---|---|---|---|---|---|---|---|
| pergolide | 2105 | 47811 | 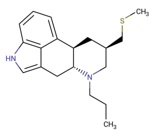 | CCCN1C[C@H](CSC)C[C@H]2[C@H]1CC1=CNC3=C1C2=CC=C3 | YEHCICAEULNIGD-MZMPZRCHSA-N | NERVOUS SYSTEM | ***TRUE*** |
| pimavanserin | 5142 | 10071196 | 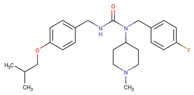 | CC(C)COC1=CC=C(CNC(=O)N(CC2=CC=C(F)C=C2)C2CCN(C)CC2)C=C1 | RKEWSXXUOLRFBX-UHFFFAOYSA-N | NERVOUS SYSTEM | ***TRUE*** |
| piribedil | 2202 | 4850 | 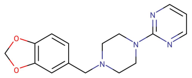 | C(N1CCN(CC1)C1=NC=CC=N1)C1=CC2=C(OCO2)C=C1 | OQDPVLVUJFGPGQ-UHFFFAOYSA-N | NERVOUS SYSTEM | ***TRUE*** |
| pramipexole | 2233 | 119570 | 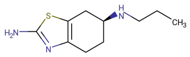 | CCCN[C@H]1CCC2=C(C1)SC(N)=N2 | FASDKYOPVNHBLU-ZETCQYMHSA-N | NERVOUS SYSTEM | ***TRUE*** |
| procyclidine | 2276 | 4919 | 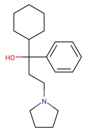 | OC(CCN1CCCC1)(C1CCCCC1)C1=CC=CC=C1 | WYDUSKDSKCASEF-UHFFFAOYSA-N | NERVOUS SYSTEM | ***TRUE*** |
| rasagiline | 3521 | 3052776 | 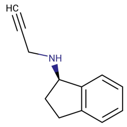 | C#CCN[C@@H]1CCC2=C1C=CC=C2 | RUOKEQAAGRXIBM-GFCCVEGCSA-N | NERVOUS SYSTEM | ***TRUE*** |
| rivastigmine | 2392 | 77991 | 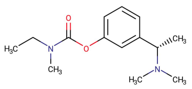 | CCN(C)C(=O)OC1=CC=CC(=C1)[C@H](C)N(C)C | XSVMFMHYUFZWBK-NSHDSACASA-N | NERVOUS SYSTEM | ***TRUE*** |
| ropinirole | 2402 | 5095 | 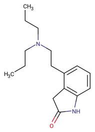 | CCCN(CCC)CCC1=CC=CC2=C1CC(=O)N2 | UHSKFQJFRQCDBE-UHFFFAOYSA-N | NERVOUS SYSTEM | ***TRUE*** |

| | | | | | | | |
|---|---|---|---|---|---|---|---|
| rotigotine | 2407 | 59227 | 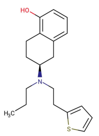 | CCCN(CCC1=CC=CS1)[C@H]1CCC2=C(O)C=CC=C2C1 | KFQYTPMOWPVWEJ-INIZCTEOSA-N | NERVOUS SYSTEM | ***TRUE*** |
| safinamide | 4921 | 131682 | 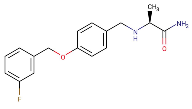 | C[C@H](NCC1=CC=C(OCC2=CC(F)=CC=C2)C=C1)C(N)=O | NEMGRZFTLSKBAP-LBPRGKRZSA-N | NERVOUS SYSTEM | ***TRUE*** |
| selegiline | 2429 | 26757 | 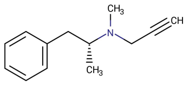 | C[C@H](CC1=CC=CC=C1)N(C)CC#C | MEZLKOACVSPNER-GFCCVEGCSA-N | NERVOUS SYSTEM | ***TRUE*** |
| tolcapone | 2697 | 4659569 | 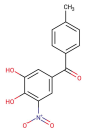 | CC1=CC=C(C=C1)C(=O)C1=CC(=C(O)C(O)=C1)[N+]([O-])=O | MIQPIUSUKVNLNT-UHFFFAOYSA-N | NERVOUS SYSTEM | ***TRUE*** |
| trihexyphenidyl | 2745 | 5572 | 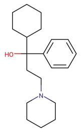 | OC(CCN1CCCCC1)(C1CCCCC1)C1=CC=CC=C1 | HWHLPVGTWGOCJO-UHFFFAOYSA-N | NERVOUS SYSTEM | ***TRUE*** |
